# A Riemannian approach to predicting brain function from the structural connectome

**DOI:** 10.1101/2021.10.27.465906

**Authors:** Oualid Benkarim, Casey Paquola, Bo-yong Park, Jessica Royer, Raúl Rodríguez-Cruces, Reinder Vos de Wael, Bratislav Misic, Gemma Piella, Boris C. Bernhardt

## Abstract

Ongoing brain function is largely determined by the underlying wiring of the brain, but the specific rules governing this relationship remain unknown. Emerging literature has suggested that functional interactions between brain regions emerge from the structural connections through mono-as well as polysynaptic mechanisms. Here, we propose a novel approach based on diffusion maps and Riemannian optimization to emulate this dynamic mechanism in the form of random walks on the structural connectome and predict functional interactions as a weighted combination of these random walks. Our proposed approach was evaluated in two different cohorts of healthy adults (Human Connectome Project, HCP; Microstructure-Informed Connectomics, MICs). Our approach outperformed existing approaches and showed that performance plateaus approximately around the third random walk. At macroscale, we found that the largest number of walks was required in nodes of the default mode and frontoparietal networks, underscoring an increasing relevance of polysynaptic communication mechanisms in transmodal cortical networks compared to primary and unimodal systems.

## Introduction

Neuroscience has increasingly embraced a paradigm shift, away from focusing on single regions towards conceptualizations that emphasize the analysis of the brain as a complex, interconnected network (Avena-Koenigsberger et al., 2018; Bassett et al., 2017; Bullmore & Sporns, 2009; Fornito et al., 2013; Sporns et al., 2005). With progress in multimodal imaging and modelling, in particular advances in diffusion MRI acquisition and tractographic reconstructions, it is now possible to visualize the structural connectome (SC), as a representation of structural wiring with increasing biological validity (Hagmann et al., 2007). Mapping the SC has critical appeal to network neuroscience, as it is generally considered to provide the physical wiring diagram of the brain that shapes and constrains ongoing brain function and dynamics, which in turn are thought to underlie the emergence of cognitive functions and behavior (Bressler & Menon, 2010).

Although a fundamental goal of systems neuroscience is to identify how structure gives rise to ongoing brain function, establishing a direct link between the SC and the functional connectome (FC) as a representation of ongoing signal interactions remains an open problem. While neural signals can be immediately transmitted between anatomically connected locations in the SC, the principles governing the flow of information between different unconnected regions of the brain remain elusive. In both humans and non-human primates, the strength of a structural connection linking two brain regions has shown to be a relatively robust predictor of the strength of their functional interaction (Honey et al., 2009; Shen et al., 2012; Skudlarski et al., 2008). Nonetheless, the predictive accuracy of the SC is far from perfect, especially when aiming to explain the functional connectivity between anatomically unconnected regions, which may be mediated by polysynaptic communication paths (Damoiseaux & Greicius, 2009; Goñi et al., 2014; Honey et al., 2009). Moreover, the strength of the structure-function relationship has generally been recognized to vary across the brain, with a tight coupling of structural and functional connectivity profiles in unimodal sensory regions that is gradually relaxed in higher-order transmodal association areas, notably regions of the default mode and frontoparietal networks (Baum et al., 2020; Vázquez-Rodríguez et al., 2019).

Several approaches have been proposed to explain the mapping between structural and functional networks, including statistical associative techniques (Mišić et al., 2016), biophysical models (Breakspear, 2017; Deco et al., 2013; Honey et al., 2009; Robinson, 2012; Wang et al., 2019), structural connectome harmonics (Abdelnour et al., 2014, 2018; Becker et al., 2018; Rosenthal et al., 2018), network communication models (Avena-Koenigsberger et al., 2018; Bazinet et al., 2021; Goñi et al., 2014; Mišić et al., 2015), and deep learning methods (Rosenthal et al., 2018; Sarwar et al., 2021). Among these approaches, those based on the eigenvectors on the SC and network communication have attracted mounting interest recently (Abdelnour et al., 2018; Atasoy et al., 2016; Becker et al., 2018; Gabay et al., 2018; Surampudi et al., 2018; Tewarie et al., 2020; Wang et al., 2017). These approaches incorporate polysynaptic communication mechanisms through more than one structural connection to account for the flow of information between not only directly connected regions but also intermediary pathways (Atasoy et al., 2016; Seguin et al., 2020; Suárez et al., 2020). By working on the structural embeddings, network communication can be modelled in a straightforward manner based on random walks on the SC (*i.e*., signal diffusion through the entire SC) or as combinations of the structural eigenvectors (Tewarie et al., 2020). As such, these techniques allow for the modelling of both mono-as well as polysynaptic communication mechanisms to incorporate increasingly high-order structural interactions, which may ultimately reconstruct a dense FC from a relatively sparse SC representation (Honey et al., 2009; Suárez et al., 2020).

Here, we propose a novel approach to predict FC from SC and better understand structure-function coupling in the human brain. The proposed approach is formulated as a kernel fusion method, where each kernel can be depicted as a putative intermediate state while the brain is propagating information through the static white matter fibers. These multi-scale diffusion kernels are implemented as random walks on the structural eigenspace, by leveraging a diffusion maps framework to identify low dimensional components describing variance in SC (Coifman et al., 2005; Coifman & Lafon, 2006). Since the functional diffusion coordinates we attempt to synthesize may have a different orientation than their analogous structural coordinates, we need a transformation to align their low-dimensional representations in manifold space. To do so, we formulate our task as a Riemannian optimization problem over the product manifold of rotations (Absil et al., 2009; Hu et al., 2020), where each rotation is used to identify the optimal paths of a specific length, and subsequently build the corresponding intermediate diffusion kernels. The proposed approach is illustrated in **Figure 1**. The workflow was evaluated on the prediction of FC from SC at the individual level rather than group level. Results are reported for two different datasets, namely: 326 unrelated subjects from the Human Connectome project (Van Essen et al., 2013) and 50 unrelated subjects from the Microstructure-Informed Connectomics dataset (Royer et al., 2021). We used the proposed approach to: *i)* study the relationship between the length of the random walks (*i.e*., length of indirect paths between brain regions) and prediction accuracy (quantified via Pearson’s correlation coefficient), *ii)* test the contribution and identify the weighting schemes to define the SC that best explain the observed brain function, *iii)* perform region- and network-specific analyses of FC prediction as a function of path length, and *iv)* compare the prediction performance of the proposed approach with several state-of-the-art methods.

**Figure 1.**
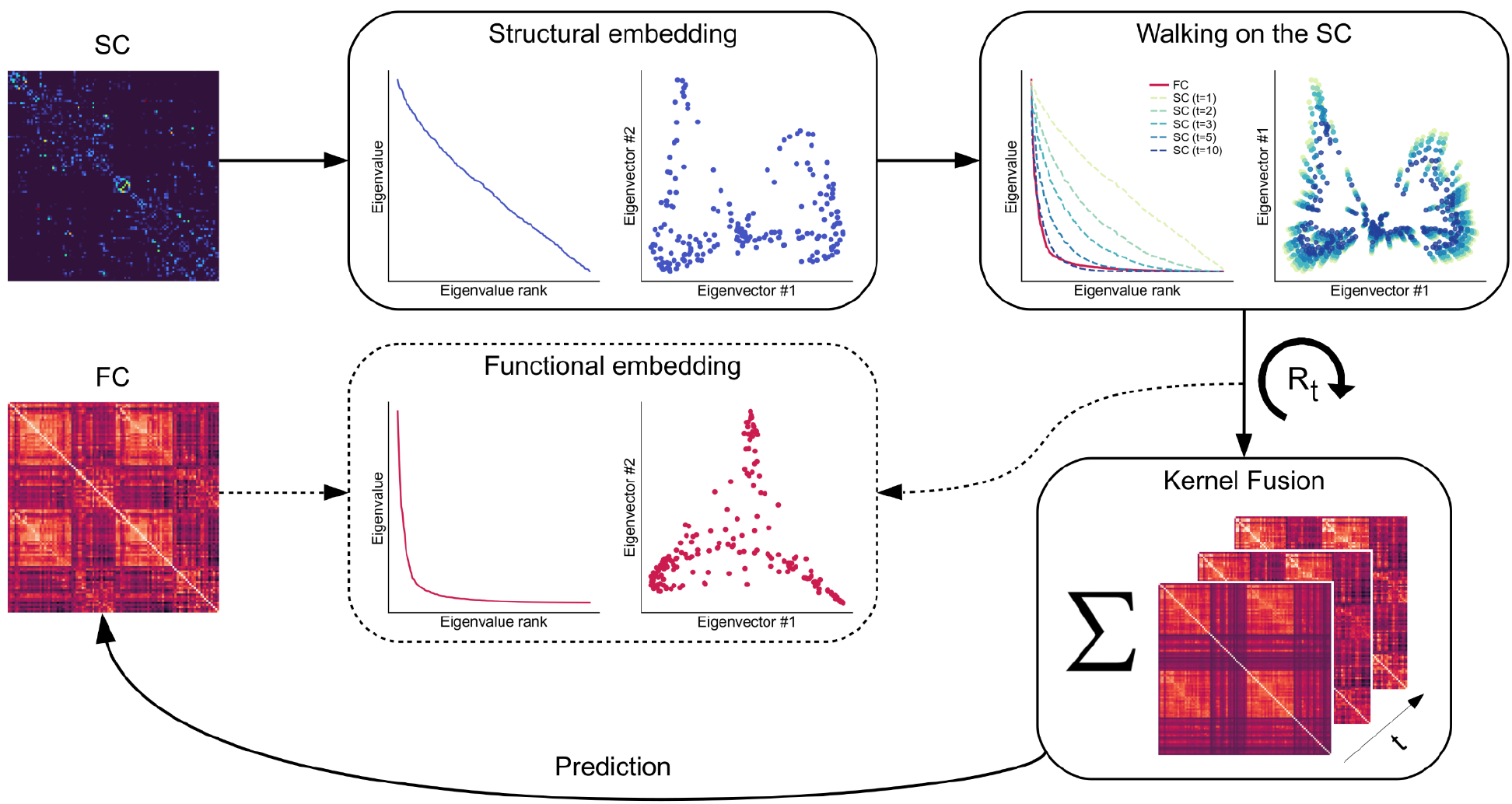
Schematic of the proposed approach. Given a pair of structural (SC) and functional (FC) connectivity matrices, first we use diffusion maps to obtain the structural embedding of the SC. The functional embedding (dashed) is only shown for visualization purposes, to illustrate the differences that might exist between the structural and functional spectra and also their eigenvectors. By increasing the diffusion time t (see Walking on the SC), we can see that the structural eigenvalues approximate the functional eigenvalues shown in red. For each diffusion time, we can obtain a different representation of the structural diffusion coordinates. The larger the diffusion time is, the closer the brain regions are to each other in the structural embedding, and hence their pairwise connectivity is increased. Then, a rotation matrix is used for each diffusion time to obtain a kernel that represents the predicted FC at time t. This can be seen as finding a rotation matrix to align to structural embedding to the functional one, see dashed line. The kernels are then fused to obtain the final predicted FC matrix.

## MATERIALS AND METHODS

### Datasets

We evaluated our proposed approach on diffusion magnetic resonance imaging (dMRI) and resting-state functional MRI (rs-fMRI) data provided by the Human Connectome Project (HCP) repository (Van Essen et al., 2013), and on the Microstructure-Informed Connectomics (MICs) dataset (Royer et al., 2021).

All MRI data used in this study were publicly available and anonymized. For HCP, we used data from the minimally processed S900 release. Participants who did not complete full imaging data and who had family relationships were excluded, resulting in a total of 326 participants (mean ± SD age = 28.56 ± 3.73 years; 55% females). Participant recruitment procedures and informed consent forms, including consent to share de-identified data, were previously approved by the Washington University Institutional Review Board as part of the HCP. For MICs, the data consist of 50 healthy volunteers (29.82 ± 5.73 years; 21 females) scanned between April 2018 and September 2020. All participants denied a history of neurological illness. The MICs dataset was approved by the Ethics Committee of the Montreal Neurological Institute and Hospital. Written informed consent, including a statement for openly sharing all data in anonymized form, was obtained from all participants.

### MRI acquisition

HCP participants were scanned using a Siemens Skyra 3T at Washington University. The T1-weighted (T1w) images were acquired using a magnetization-prepared rapid gradient echo (MPRAGE) sequence (repetition time (TR) = 2,400 ms; echo time (TE) = 2.14 ms; field of view (FOV) = 224 × 224 mm^2^; voxel size = 0.7 mm^3^; and number of slices = 256). The T2-weighted (T2w) structural data were obtained with the T2-SPACE sequence, with an identical geometry as the T1w data but different TR (3,200 ms) and TE (565 ms). The dMRI data were acquired with the spin-echo echo-planar imaging (EPI) sequence (TR = 5,520 ms; TE = 89.5 ms; FOV = 210 × 180 mm^2^; voxel size = 1.25 mm^3^; b-value = three different shells *i.e*., 1,000, 2,000, and 3,000 s/mm^2^; number of diffusion directions = 270; and number of b0 images = 18). The rs-fMRI data were collected using a gradient-echo EPI sequence (TR = 720 ms; TE = 33.1 ms; FOV = 208 × 180 mm^2^; voxel size = 2 mm^3^; number of slices = 72; and number of volumes = 1,200). During the rs-fMRI scan, participants were instructed to keep their eyes open looking at a fixation cross. Two sessions of rs-fMRI data were acquired; each of them contained data of left-to-right and right-to-left phase-encoded directions, providing up to four time series per participant.

For MICs, participants were scanned at the McConnell Brain Imaging Centre of the Montreal Neurological Institute and Hospital on a 3T Siemens Magnetom Prisma-Fit equipped with a 64-channel head coil. Participants underwent a T1w structural scan, followed by multi-shell dMRI and rs-fMRI. In addition, a pair of spin-echo images was acquired for distortion correction of individual rs-fMRI scans. Two T1w scans with identical parameters were acquired with a 3D-MPRAGE sequence (TR = 2300 ms, TE = 3.14 ms, TI = 900 ms, flip angle = 9°, iPAT = 2, partial Fourier = 6/8, voxel size = 0.8 mm^3^, matrix = 320×320, and number of slices = 224). Both T1w scans were visually inspected to ensure minimal head motion before they were submitted to further processing. A spin-echo EPI sequence with multi-band acceleration was used to obtain dMRI data, consisting of three shells with b-values 300, 700, and 2000s/mm2 and 10, 40, and 90 diffusion weighting directions per shell, respectively (TR = 3500 ms, TE = 64.40 ms, voxel size = 1.6 mm^3^, flip angle = 90°, refocusing flip angle = 180°, FOV = 224×224 mm^2^, slice thickness = 1.6 mm, mb factor = 3, echo spacing = 0.76 ms, number of b0 images = 3). One rs-fMRI scan was acquired using multiband accelerated 2D-BOLD EPI (TR = 600 ms, TE = 30 ms, voxel size = 3mm^3^, flip angle = 52°, FOV = 240×240 mm^2^, slice thickness = 3 mm, mb factor = 6, echo spacing = 0.54 ms). Participants were instructed to keep their eyes open, look at a fixation cross, and not fall asleep. A complete list of acquisition parameters can be found in the detailed imaging protocol provided by (Royer et al., 2021).

### Data preprocessing

HCP data underwent the initiative’s minimal preprocessing pipelines (Glasser et al., 2013). In brief, structural MRI data underwent gradient nonlinearity and b0 distortion correction, followed by co-registration between the T1w and T2w data using a rigid-body transformation. Bias field correction was performed by capitalizing on the inverse intensities from the T1- and T2-weighting. Processed data were nonlinearly registered to MNI152 space and the white and pial surfaces were generated by following the boundaries between different tissues (Dale et al., 1999; Fischl, 2012; Fischl et al., 1999a, 1999b). The white and pial surfaces were averaged to generate a mid-thickness surface, which was used to generate the inflated surface. The spherical surface was registered to the Conte69 template with 164k vertices (Van Essen et al., 2012) using MSMAll (Glasser et al., 2016; Robinson et al., 2014) and downsampled to a 32k vertex mesh. The dMRI data underwent b0 intensity normalization, and EPI distortions were corrected by leveraging reversed phase-encoded directions. The dMRI data was also corrected for eddy current distortions and head motion. The rs-fMRI data preprocessing involved corrections for EPI distortions and head motion, and fMRI data were registered to the T1w data and subsequently to MNI152 space. Magnetic field bias correction, skull removal, and intensity normalization were performed. Noise components attributed to head movement, white matter, cardiac pulsation, arterial, and large vein related contributions were automatically removed using FIX (Salimi-Khorshidi et al., 2014). The minimal preprocessing with FIX-denoising pipeline of the HCP performs a high-pass filtering with a cutoff of 2,000 s full width at half maximum (FWHM) (Glasser et al., 2013). Preprocessed time series were mapped to standard grayordinate space, with a cortical ribbon-constrained volume-to-surface mapping algorithm. The total mean of the time series of each left-to-right/right-to-left phase-encoded data was subtracted to adjust the discontinuity between the two datasets and they were concatenated to form a single time series data.

For the MICs dataset, T1w images were anonymized and de-identified by defacing all structural volumes. Each T1w scan was deobliqued and reoriented. T1w scans were then linearly co-registered and averaged, automatically corrected for intensity nonuniformity (Tustison et al., 2010), and intensity normalized. Resulting images were skull-stripped, and subcortical structures were segmented using FSL FIRST (Jenkinson et al., 2012). Cortical surface segmentations were generated from native T1w scans using FreeSurfer 6.0 (Dale et al., 1999; Fischl, 2012; Fischl et al., 1999a, 1999b). The dMRI data were preprocessed using MRtrix (Tournier et al., 2012, 2019). The dMRI data underwent b0 intensity normalization, and were corrected for susceptibility distortion, head motion, and eddy currents. Required anatomical features for tractography processing were co-registered to native dMRI space using affine transformation tools implemented in Advanced Neuroimaging Tools (ANTs) (Tustison & Avants, 2013). Diffusion processing and tractography were performed in native dMRI space. For rs-fMRI, images were pre-processed using AFNI (Cox, 1996) and FSL (Jenkinson et al., 2012). The first five volumes were discarded to ensure magnetic field saturation. Images were reoriented, and motion as well as distortion corrected. Nuisance variable signal was removed using ICA-FIX (Salimi-Khorshidi et al., 2014) and by performing spike regression. Volume timeseries were averaged for registration to native FreeSurfer space using boundary-based registration (Greve & Fischl, 2009). Native timeseries were mapped to individual surface models using a boundary-based registration and smoothed using a Gaussian kernel (FWHM=10mm, smoothing performed on native midsurface mesh) using workbench. The preprocessing of the MICs dataset was performed with micapipe (https://micapipe.readthedocs.io).

### Functional and structural connectome generation

To estimate FC matrices, individual rs-fMRI timeseries mapped to individual surface models were averaged within parcels defined by the cortical parcellation scheme. Cortical timeseries were sampled from each vertex of the native FreeSurfer space cortical surface segmentation, and averaged within surface parcels. Individual functional connectomes were generated by cross-correlating all nodal timeseries.

SC representations were generated from preprocessed dMRI data using MRtrix (Tournier et al., 2012, 2019). Different tissue types of cortical and subcortical grey matter, white matter, and cerebrospinal fluid were segmented using T1-weighted image for anatomical constrained tractography (Smith et al., 2012). Multi-shell and multi-tissue response functions were estimated (Christiaens et al., 2015) and constrained spherical-deconvolution and intensity normalization were performed (Jeurissen et al., 2014). The initial tractogram was generated with 40 million streamlines, with a maximum tract length of 250 and a fractional anisotropy cutoff of 0.06. Spherical-deconvolution informed filtering of tractograms (SIFT2) was applied to reconstruct whole brain streamlines weighted by cross-section multipliers (Smith et al., 2015). To build a structural connectome, the reconstructed cross-section streamlines were mapped to selected cortical parcellation schemes.

### Proposed framework

Let 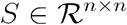 be the connectivity matrix representing a given SC, where each entry *S*(*i*, *j*) is the weight of the edge connecting the *i*-th and *j*-th cortical locations, computed as the total number of streamlines connecting both locations, such that *S*(*i, j*)=*S*( *j*, *i*) and *S*(*i*, *j*)≥ 0,∀ *i, j*=1,…,*n*. Our purpose is to predict the FC, the correlations of resting-state functional signals, from its corresponding SC (i.e., *S*). To do so, we leverage diffusion maps (Coifman & Lafon, 2006). We first proceed by normalizing the SC matrix to define the diffusion operator *P*:

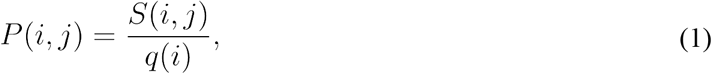

where 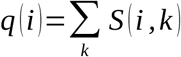 denotes the degree in the connectome, such that 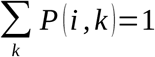. Now *P_ij_* can be viewed as the probability for a random walker on the SC *S* to make a step from the *i*-th to *j*-th cortical locations. As *P* is not symmetric, we can further define a symmetric operator Δ:

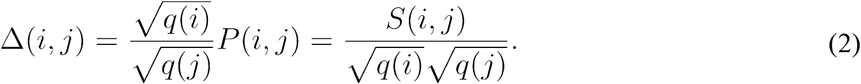

In matrix notation, we have that *Δ*=Q^-1/2^ SQ^-1/2^ where *Q* denotes the degree matrix of *S* (a diagonal matrix such that 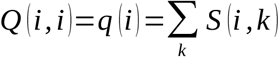. Using spectral theory, it can be shown that Δ has the following eigendecomposition:

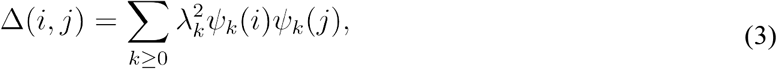

where 1 = *λ*_0_≥∨*λ*_1_∨≥∨*λ*_2_∨≥… is the eigenspectrum and {*ψ_k_*} the corresponding eigenvectors of *Δ*. This operator shares the same spectrum with *P* and its eigenvectors are orthogonal, unlike those of *P* (Coifman & Hirn, 2014). Now, a walk of length *t* in the SC can be represented by the diffusion maps *Ψ_t_* as follows:

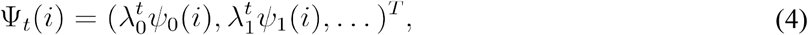

where *T* stands for transpose. A desirable property of the diffusion map *Ψ_t_* is that it embeds the data into a Euclidean space in which the Euclidean distance is equal to the diffusion distance *D_t_*:

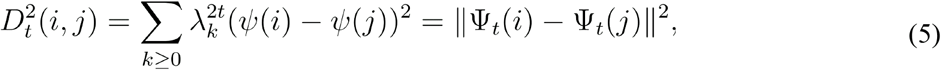

where ‖·‖ denotes the *l*_2_-norm. With this metric, we can capture the connectivity of two cortical locations for each length of the walks *t* in the SC. Note that as *t* grows, the diffusion distance between the cortical locations will decrease and will be mainly driven by the first diffusion coordinates, i.e., those coordinates corresponding to the largest eigenvalues. This will allow us to approximate the distances based solely on the dominant eigenvectors and reduce the dimensionality of the diffusion maps.

Let 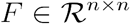 be the FC matrix, built using time-series correlation analysis of an fMRI scan from the same subject. Since diffusion maps define a Euclidean space, we propose to predict the FC using a kernel fusion approach:

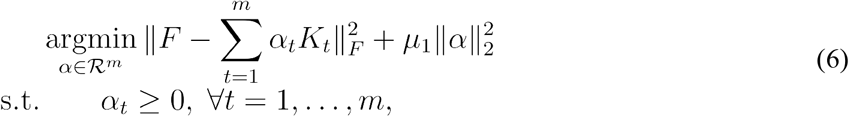

where 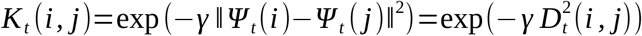 is a radial basis function (RBF) kernel built from the diffusion maps *Ψ_t_* for each diffusion time *t*, and *γ* is the kernel bandwidth; *m* is the total number of walks considered, *α_t_*≥0 is the coefficient corresponding to the RBF kernel *K_t_*, *μ*_1_ is a tradeoff parameter, and ‖ · ‖_*F*_ is the Frobenius norm. Given that *K_t_* (*i*, *j*) ∈ [0, 1] and *F*(*i, j*)∈[−1, 1], in our setting we scale the FC matrices to the range [0, 1] for training and undo this operation after prediction. Since we will assess performance using Pearson’s correlation coefficient, which is invariant under positive linear transformations, this scaling has no effect on the results.

Although we can approximate the functional eigenvalues with an increasing number of walks in the structural embedding, as shown in **Figure 1**, the structural and functional embeddings do not share the same diffusion coordinates. We therefore propose to find a transformation of the structural embedding to more faithfully reconstruct the FC. Let 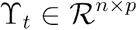 be a matrix representing the diffusion coordinates of the SC at time *t* and *p*≤*n*, we aim to find a rotation matrix 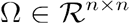 to transform the diffusion coordinates as follows:

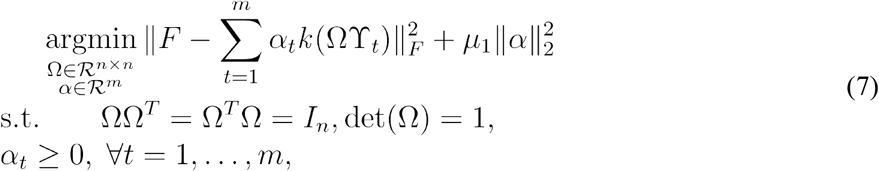

where *I_n_* is an *n*×*n* identity matrix, *det* (·) stands for matrix determinant, and ¿ and *Γ_t_* = *ΩY_t_*, with *Γ_t_* denoting the rotated structural diffusion map, and *Γ_t_* (*i*) the *i*-th row of *Γ_t_*. It is worth noting that the scaling of the eigenvectors is different for each diffusion time *t*, and because of the spectrum decay, we will be having fewer and fewer diffusion coordinates contributing to the computation of the RBF kernels. This will be producing a different kernel *K_t_* for each diffusion time *t*. Based on this observation, we extend our approach to include a rotation for each diffusion time:

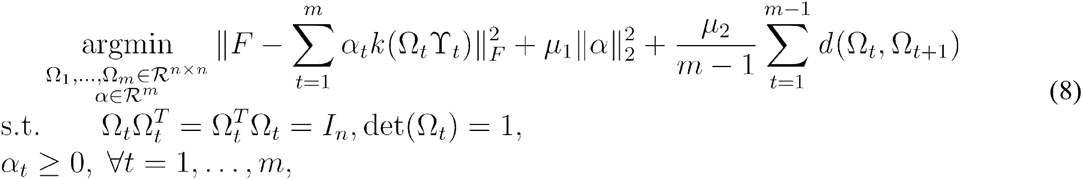

where we include a third term and its trade-off parameter *μ_2_* to avoid overfitting, with *d* (*Ω_t_,Ω*_*t*+1_) denoting the distance between two consecutive rotation matrices. When *μ*_2_=0, no restrictions are imposed on the rotation matrices, whereas when *μ_2_*> 0, this term enforces the rotations of consecutive diffusion times to be similar to each other. In the extreme case where *μ_2_* tends to infinity, the problem amounts to finding one single rotation shared by all diffusion times (*as shown in* Eq. (7)). Biologically, one can think of these rotation matrices as identifying the optimal paths through which to propagate information between different regions of the brain. Since we have one rotation matrix for each diffusion time, each rotation will identify paths of a specific length. That is, the rotation corresponding to diffusion time *t* will attempt to find the optimal paths of length *t* that connect a pair of brain regions.

To infer the latent variables in our problem, we employ an alternating optimization technique. We minimize the cost function in Eq. (8) for each output variable, while holding the estimates of the other unknowns constant. Note that, for brevity, in our formulation we only considered a single subject, but the optimization is performed for multiple subjects. To determine the set of rotations, we recast our cost function to a Riemannian manifold optimization problem (Absil et al., 2009; Hu et al., 2020). Riemannian optimization translates a constrained optimization problem into an unconstrained optimization problem where the constraints are implicitly defined by the search space. Particularly, for our problem, the target manifold is the Special Orthogonal group 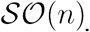 For more than one rotation, we can define the target manifold as the product manifold of rotations, 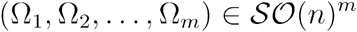:

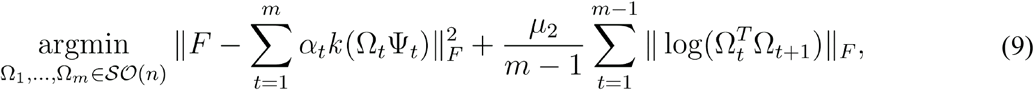

where the distance between rotation matrices 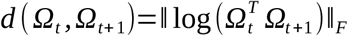 is suitably chosen to be the geodesic distance on 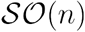 and log(·) denotes the matrix logarithm (Boumal et al., 2014). Given that kernels corresponding to consecutive numbers of walks are similar, this term imposes that their rotation matrices must lie close to each other on the rotation manifold. The problem in Eq. (9) can be solved using Riemannian optimization algorithms. In our case, we use the Riemannian conjugate gradient algorithm (Absil et al., 2009) as implemented in Pymanopt (Townsend et al., 2016).

Once we have estimated the rotation matrices, the next step is solving for *α*, which amounts to ridge regression with non-negativity constraints on the coefficients. Let *X* =(*uvec*(*K_1_*),*uvec*(*K_2_*),…,*uvec*(*K_m_*)) and *y*=*uvec*(*F*), where *uvec*(·) returns the upper triangular part of a symmetric matrix, without the elements in the main diagonal. This optimization problem amounts to:

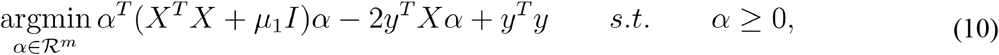

where the third term does not affect the optimization. This problem can be solved with conventional quadratic programming. The proposed approach is implemented in Python and the code is publicly available at (https://github.com/MICA-MNI/micaopen/sf_prediction).

### Experimental settings

The performance of the proposed approach in predicting FC from SC was assessed in the HCP and MICs datasets. For MICs, prediction accuracy was based on a 3-fold cross-validation (CV) strategy based on all 50 individuals in the dataset. For HCP (n=326), the data was randomly split in 3 subsets: 50 individuals were selected for CV, 250 for holdout, and the remaining 26 individuals were used for parameter tuning. Following previous studies, prediction performance was reported in terms of Pearson’s correlation coefficient based on the upper triangular parts (excluding the main diagonal) of both empirical and predicted FC matrices. Regarding parameter tuning, we used an independent subset (26 subjects) from the HCP dataset to find the optimal values for the hyperparameters (i.e., *μ*_1_, *μ*_2_, and *γ* of the RBF kernels) of our proposed approach. Note that this subset was not used to report any results. The best values for both *μ*_1_and *μ*_2_ were chosen from a grid of 9 equidistant points in logarithmic scale in the interval [1e-4, 1e4]. The optimal value for *μ*_1_ was found to be 100. For the Riemannian regularization, our initial benchmarking showed a negligible improvement for *μ*_2_*≤*0.001, and we therefore chose to report our results for *μ*_2_= 0. For the RBF kernel, *γ* was chosen to be the standard deviation of the diffusion distances for each random walk. These optimal hyperparameter values were then used to assess performance in both the HCP and MICs datasets.

Our proposed approach was tested using SC and FC matrices built based on two different cortical atlases: *i)* a parcellation derived using functional MRI data (Schaefer et al., 2018), and *ii)* a structurally-defined parcellation based on a more fine-grained clustering of the well-established Desikan Killiany parcellation (Desikan et al., 2006; Vos de Wael et al., 2020). To assess the robustness of the proposed approach across different spatial scales, our experiments were repeated for two different parcellation granularities, using parcellations with 100 and 200 cortical regions. We also explored different weighting schemes to define the SC matrices in explaining ongoing brain function. We analyzed three versions of the SC: *i)* binary SC, where each entry in the SC matrix is set to 1 only if there is at least one streamline connecting the corresponding regions of the brain, *ii)* length-based SC, which is built using an RBF kernel based on the mean length of the streamlines connecting pairs of brain regions, and *iii)* count-based SC, which is defined based in the number of streamlines connecting each pairs of brain regions. Unless otherwise stated, our experiments are based on SC matrices defined using the latter weighting scheme (*i.e*., count-based). We also investigated the contribution of diffusion time at network level, and the role it plays in strengthening the structure-function coupling and its relationship with the principal functional gradient (Margulies et al., 2016). The principal functional gradient was estimated using BrainSpace (Vos de Wael et al., 2020).

We compared the performance of the proposed approach with state-of-the-art methods. Among those based on the structural eigenvectors, we compared our approach to the Multiple Kernel Learning (MKL) (Surampudi et al., 2018) and Spectral (Becker et al., 2018) approaches. Briefly, in the MKL approach, multiple diffusion kernels on the SC are linearly combined using LASSO (Tibshirani, 1996) to predict FC. The Spectral approach, on the other hand, learns a shared functional embedding and a mapping from the functional to the structural eigenvalues for each individual, which are then used to build the predicted FC matrices. Note that, in the Spectral approach, the individual structural eigenvectors are not taken into consideration to predict FC. Outside the eigenvector-based category, we included results using the single Laplacian-based diffusion kernel (SDK) proposed in (Abdelnour et al., 2014), and the series expansion approach (NLSA) in (Meier et al., 2016). SDK defines a single diffusion kernel at a specific scale (or diffusion time) from the symmetric normalized Laplacian matrix of the SC. In our work, the optimal scale was chosen so that it provided the best performance in the training data. NLSA, on the other hand, predicts FC as a truncated Taylor series expansion of the SC (Tewarie et al., 2020). We used code provided by the authors for both MKL (https://github.com/govindasurampudi/MKL) and Spectral (https://brainopt.github.io/spectral-mapping). Finally, our approach used multiple kernels along with their corresponding rotation matrices. To elucidate the contribution of each of these components, we included two additional versions of our method to the comparison table: 1) “SingleLength” only used one single kernel or random walk for prediction, and 2) “SharedRot” used a single rotation matrix shared by all the random walks.

**Table 1.** Comparison of functional connectivity prediction accuracy with state-of-the-art methods. Performance is reported using the average Pearson’s correlation coefficient (and standard deviation) between the upper triangular parts of empirical and predicted FC matrices (excluding the main diagonal) using SDK, NLSA, MKL, Spectral and the 3 versions of our proposed approach: *i)* consider random walks of a specific length (SingleLength), *ii)* one single rotation shared by all random walks (SharedRot), and *iii)* one rotation for each length of the random walks (Proposed). Comparisons of prediction accuracy were carried out for 100- and 200-node functional and structural parcellations. Results are reported for a 3-fold cross validation (CV) and in the holdout data in HCP. For NLSA, Spectral and all the versions of our proposed approach, reported performances correspond to using random walks of length 10, and 16 for MKL. For SDK, the diffusion time was chosen to be the one that achieved the best performance in the training set. For findings in the MICs dataset, see Table S1.

|  |  | Functional |  | Structural |  |
| --- | --- | --- | --- | --- | --- |
|  |  | CV | Holdout | CV | Holdout |
| 100 | SDK | 0.238±0.029 | 0.234±0.027 | 0.343±0.057 | 0.347±0.069 |
|  | NLSA | 0.210±0.037 | 0.216±0.036 | 0.351±0.050 | 0.348±0.061 |
|  | MKL | 0.713±0.101 | 0.726±0.106 | 0.664±0.110 | 0.705±0.121 |
|  | Spectral | 0.779±0.059 | 0.761±0.082 | 0.703±0.090 | 0.669±0.105 |
|  | SingleLength | 0.757±0.050 | 0.767±0.051 | 0.731±0.052 | 0.748±0.050 |
|  | SharedRot | 0.733±0.050 | 0.751±0.048 | 0.701±0.068 | 0.728±0.056 |
|  | Proposed | **0.802±0.054** | **0.804±0.060** | **0.751±0.065** | **0.764±0.054** |
| 200 | SDK | 0.241±0.027 | 0.237±0.025 | 0.245±0.043 | 0.243±0.035 |
|  | NLSA | 0.211±0.039 | 0.213±0.030 | 0.241±0.039 | 0.240±0.043 |
|  | MKL | 0.702±0.105 | 0.701±0.099 | 0.612±0.124 | 0.627±0.132 |
|  | Spectral | 0.741±0.073 | 0.725±0.081 | 0.681±0.083 | 0.650±0.145 |
|  | SingleLength | 0.727±0.048 | 0.738±0.044 | 0.692±0.063 | 0.700±0.054 |
|  | SharedRot | 0.701±0.056 | 0.720±0.047 | 0.539±0.112 | 0.571±0.102 |
|  | Proposed | **0.775±0.049** | **0.776±0.052** | **0.715±0.071** | **0.726±0.055** |

## RESULTS

### Walking on the structural connectome increases prediction accuracy

To faithfully reconstruct the FC from the sparse SC, we first investigated the role of diffusion time (or path length) in the prediction accuracy of our proposed approach. We explored the contribution of random walks of different lengths, ranging from 1 to 10, such that for a given maximum length of 5 for example, we considered random walks of any length below or equal to 5. In this way, we are not only including the static direct connections present in the SC, but also allow our method to incorporate information about the SC at multiple scales, *i.e*., after propagating information between indirectly connected brain regions.

The prediction accuracy of our proposed approach was assessed in two different datasets, namely: HCP and MICs. For HCP, **Figure 2a** shows boxplots with the correlation between the empirical and predicted FC matrices using structural random walks with maximum lengths ranging from 1 to 10. Results are reported in both cross-validation (using a 3-fold CV based on 50 randomly chosen subjects) and holdout data (250 subjects) subsets, and using different cortical parcellation atlases (defined using structural and functional information) and number of regions (100 and 200 parcels). These results showed that the highest performance (in terms of Pearson’s correlation coefficient) was achieved with approximately a maximum path length of 3-4. After that, performance plateaued and the change in prediction accuracy, although positive, was in most scenarios negligible. Overall, these trends were consistent in both cross-validation and holdout datasets, and across the different parcellations and numbers of regions. In the holdout set of HCP, for example, prediction accuracy improved substantially when considering random walks of length up to 3 (0.174/0.165 increase in mean Pearson’s correlation coefficient for 100/200 regions functional parcellations, and 0.085/0.112 for the analogous structural parcellations), whereas with the incorporation of additional walks (those with path lengths from 4 to 10), we only found very small improvements in performance (0.017/0.007 and 0.013/0.002 average increase for the 100/200-node functional and structural parcellations, respectively). Moreover, when comparing the type and size of the parcellations, prediction accuracy was consistently lower in structural than in functional parcellations, and dropped when increasing the number of parcels used to build the connectivity matrices. Results on the MICs dataset, based on a 3-fold crossvalidation, are shown in **Figure S1** of the supplementary materials. Similar to HCP, highest performance was achieved with random walks of length 3 or shorter, with prediction accuracy decaying when the number of cortical parcels increased. However, no clear trend was found when comparing performance based on the structural and functional parcellation (see *Comparison with state-of-the-art methods* section).

**Figure 2.**
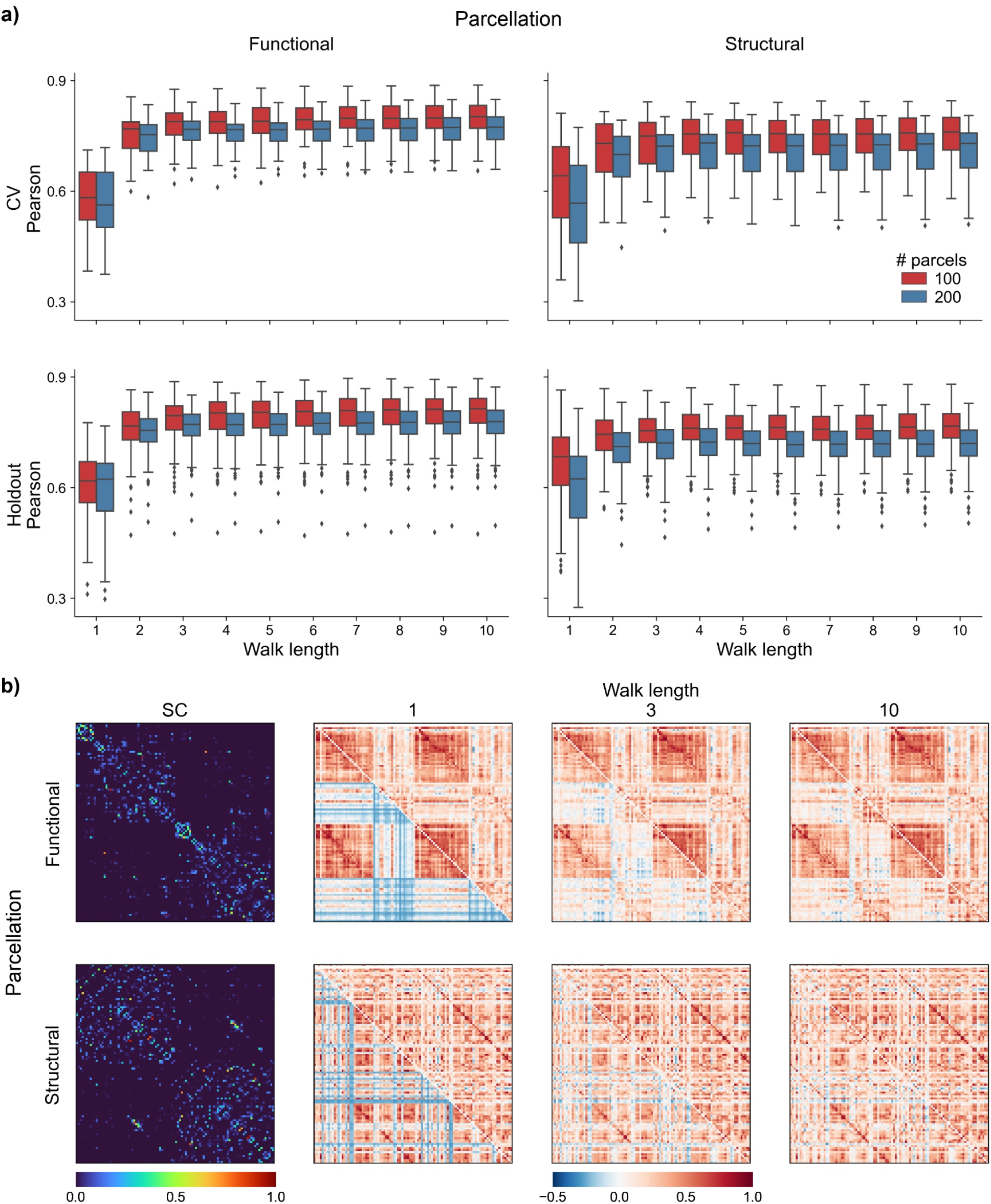
Functional connectivity prediction accuracy using random walks of different lengths in the structural connectome. **a)** Boxplots of Pearson’s correlation coefficient between empirical and estimated FC matrices using random walks of maximum length ranging from 1 to 10. Results are reported in cross-validation (3-fold CV based on 50 subjects) and holdout (250 subjects) data of HCP using functionally- (*left*) and structurally-derived cortical parcellations (*right*) with 100 and 200 regions. Boxes denote the interquartile range (IQR) between the first and third quartiles, and the line inside denotes the median. Whiskers extend to points that lie within 1.5 IQRs of the lower and upper quartiles, and the black diamonds denote outliers. **b)** Estimated functional connectivity matrices corresponding to the subjects that achieved the median Pearson’s correlation coefficient based on functional (*top*) and structural (*bottom*) parcellations with 100 parcels. From left to right: structural connectome and estimated FC matrices for random walks of length 1, 3 and 10. Empirical and estimated functional connectivity matrices are shown in upper and lower triangular parts, respectively.

To further illustrate how brain function emanates from the SC as the length of the random walks increases (*i.e*., as we increasingly incorporate indirect connections between cortical regions), **Figure 2b** displays the original SC matrices and the predicted FC matrices corresponding to the individuals that achieved the median Pearson’s correlation coefficient in the holdout subset of the HCP dataset. The predicted FC matrices corresponding to random walks of length 1, 3 and 10 are displayed in the lower triangular parts, with their respective empirical FC shown in the upper triangular parts. Results shown in **Figure 2b** are based on both the structural and functional parcellations with 100 regions. For the 200-node parcellations, results are shown in **Figure S2**. With both parcellation types and sizes, we found some similarities emerge with random walks of length 1, but these considerably increased as larger walks (indirect paths) were incorporated. This points to communication between different cortical regions through indirect paths of lengths larger than one, which highlights the importance of polysynaptic mechanisms in the emergence of the brain’s function and, at the same time, shows that only a small number of hops (path lengths ≤3) in the SC may be required to accurately predict FC, as shown by the results in **Figure 1a**. Furthermore, we also found that functionally-defined cortical parcellations were more suitable than structurally-derived ones in the prediction of FC from SC.

### Comparison of structural connectome characteristics

The structural connectivity matrices used so far in our experiments were derived from fiber density estimates, such that each entry in the matrix denotes the number of streamlines connecting the specific pair of brain regions. Here, we sought to investigate the contribution of different weighting schemes to define SC. First, we used different versions of the structural connectivity matrices: *i)* the original SC matrices, where edges carried information about the streamline count, *ii)* binary SC, where we preserved the same edges but ignored edge information (with the weights of all edges set to 1), and *iii)* length-based SC, with the same edges but the weights were based on the inverse of the average length of the streamlines. We repeated the experiments to learn the set of rotations and kernel coefficients for each version of the structural connectome.

As shown in **Figure 3a**, based on results on the holdout data of HCP and 200-node parcellations, when using SC based on streamline count we achieved the best accuracy in predicting FC, with a considerable improvement over binary SC, which in turn outperformed length-based SC. More importantly, diffusion time did not seem to have the same contribution when using binary or length-based SC matrices for prediction. With the functional parcellation, a path length of 1 with the binary SC provided better FC predictions than the count-based SC, however, as the lengths of the paths increased, the boost in performance only occurred with count-based SC. With the structural parcellation, this was even more evident, with prediction accuracy remaining almost constant as diffusion time increased when using binary and length-based SC. Following the diffusion maps framework, with increasing diffusion time, pairs of brain regions will increase the strengths of their connections according to the strengths of their immediate direct connections and those of their neighbors. For the purposes of brain function, this may suggest that the more important the ‘structural’ connection between a pair of regions is, the more fibers the brain invests in its construction. Examples of predicted FC matrices using binary and length-based SC are displayed in **Figure 3b**. These examples correspond to the subjects that achieved the mean Pearson’s correlation coefficient in the holdout set of HCP for the 200-node functional and structural parcellations. As we can see, the proposed approach produced the least accurate FC matrices when using length-based SC matrices. The fact that better prediction accuracies were achieved by binary SC matrices, which may be an oversimplified representation of the connectome, indicates that the length of a given connection (quantified as the mean length along all interconnecting streamlines) may not be as informative as its density.

**Figure 3.**
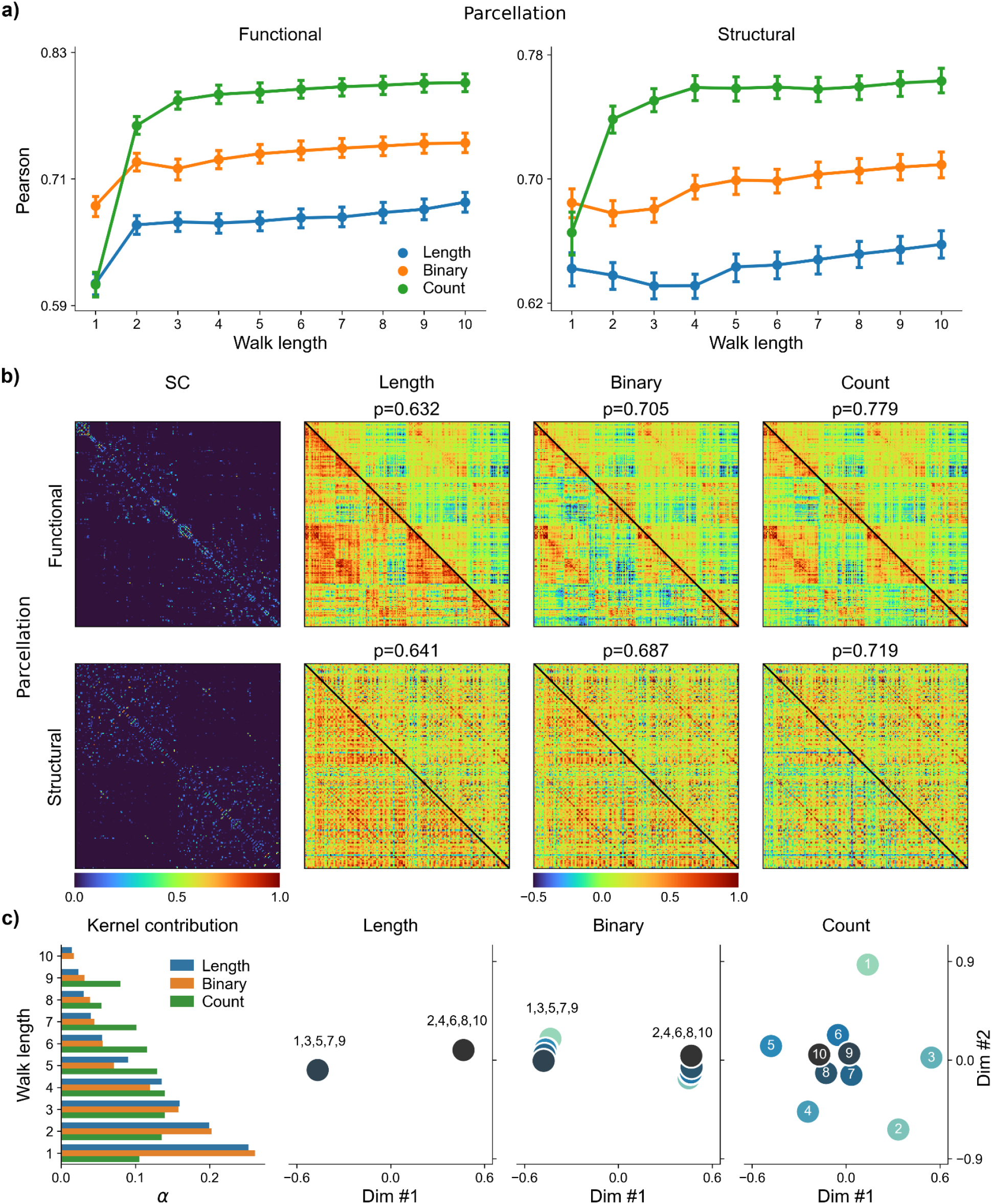
Comparison of prediction performance and model parameters based on binary, length- and count-based SC matrices. **a)** Mean Pearson correlation (and 95% confidence interval) between empirical and estimated FC matrices based on SC matrices built using streamline length, streamline count, and the binarized connectome. Results are reported for random walks of length 1 to 10 on the SC for both the 200-node functional (*left*) and structural (*right*) parcellations. **b)** Estimated FC matrices corresponding to the subjects that achieved the median Pearson’s correlation coefficient based on functional (*top*) and structural (*bottom*) parcellations with 200 parcels. From left to right: count-based SC matrices, and empirical and estimated FC matrices based on streamline length, binary SC, and streamline count. The displayed estimated connectivity matrices correspond to using random walks of length ≤10. **c)** Optimal kernel coefficients (normalized), α, and rotations matrices obtained when trained with binary, length- and count-based SC matrices using random walks of length ≤10 and the 200-node functional parcellation. Rotation scatterplots are based on the two first dimensions of multidimensional scaling (using the geodesic distance between rotation matrices). Results are reported in the holdout data of the HCP dataset.

Finally, in **Figure 3c** we can find the optimal kernel coefficients and rotations matrices learnt by our proposed approach when trained with each of the three different versions of the SC matrices. When using count-based SC, the kernels that most contributed to the final prediction corresponded to diffusion times 2-5, followed by 6 and 1. Kernels with higher diffusion times (t>6) had the least contribution. On the other hand, with both binary and length-based SC, kernel contributions decayed monotonically with increasing diffusion time. The most relevant differences, however, were found when comparing the rotation matrices learnt from the different SC versions. The rotation matrices, displayed using multidimensional scaling (based on the geodesic distance between rotations, as defined in Eq. (9)), were clustered in two groups of very similar rotations when using binary and length-based SC. On the other hand, with count-based SC matrices, we found different rotations for each diffusion time until 5, while rotations were very similar for diffusion times of 6 and greater. These results indicate that diffusion time (*i.e*., considering increasingly larger paths) is of substantial added value to the prediction of FC when using count-based SC, as opposed to binary and length-based SC data.

### Region- and network-wise analysis of predicted functional connectivity

In this section, we scrutinized our results to further investigate the role of diffusion time at the regional and network-level. **Figure 4a** shows spatial maps of prediction error (measured as *log(1-Pearson))* for different diffusion times (*i.e*., 1, 2, 3 and 10). Here we display the average prediction error in the holdout data of HCP for each cortical parcel (*i.e*., derived from row-wise correlations). There were clear improvements across the whole cortex in the quality of the predictions as increasingly longer paths were considered, although they became subtler with higher diffusion times. The cortical regions where our approach produced the largest prediction errors (irrespective of diffusion time) were confined bilaterally to the lateral temporal lobe and frontal cortices. At the network level, derived using a previous cortical decomposition into seven intrinsic functional networks (Yeo et al., 2011), prediction error decreased substantially from walk lengths of 1 to 3 (see **Figure 4b**). With higher walk lengths, we can only observe minor changes in prediction error across all networks. These findings were consistent across different parcellation granularities. Moreover, from these results we could identify two different groups of networks according to their prediction errors. FC prediction was more accurate in visual, somatomotor and both attention networks than in the default mode, frontoparietal and limbic networks. The latter were the networks that most benefited from incorporating indirect paths to the prediction, indicating that polysynaptic connections may have an important contribution to functional connectivity patterns, especially in these transmodal i.e., heteromodal and paralimbic systems. This was further confirmed when analyzing the relationships between the principal functional gradient (see **Figure 4c**) and the spatial maps of prediction error. As shown in **Figure 4d**, these maps showed very high correlations with the principal functional gradient when using short paths (Spearman’s r=0.843/0.835 in 100/200-node functional parcellation when diffusion time=1), but considerably decrease as diffusion time increased (Spearman’s r=0.250/0.338 when diffusion time=4).

**Figure 4.**
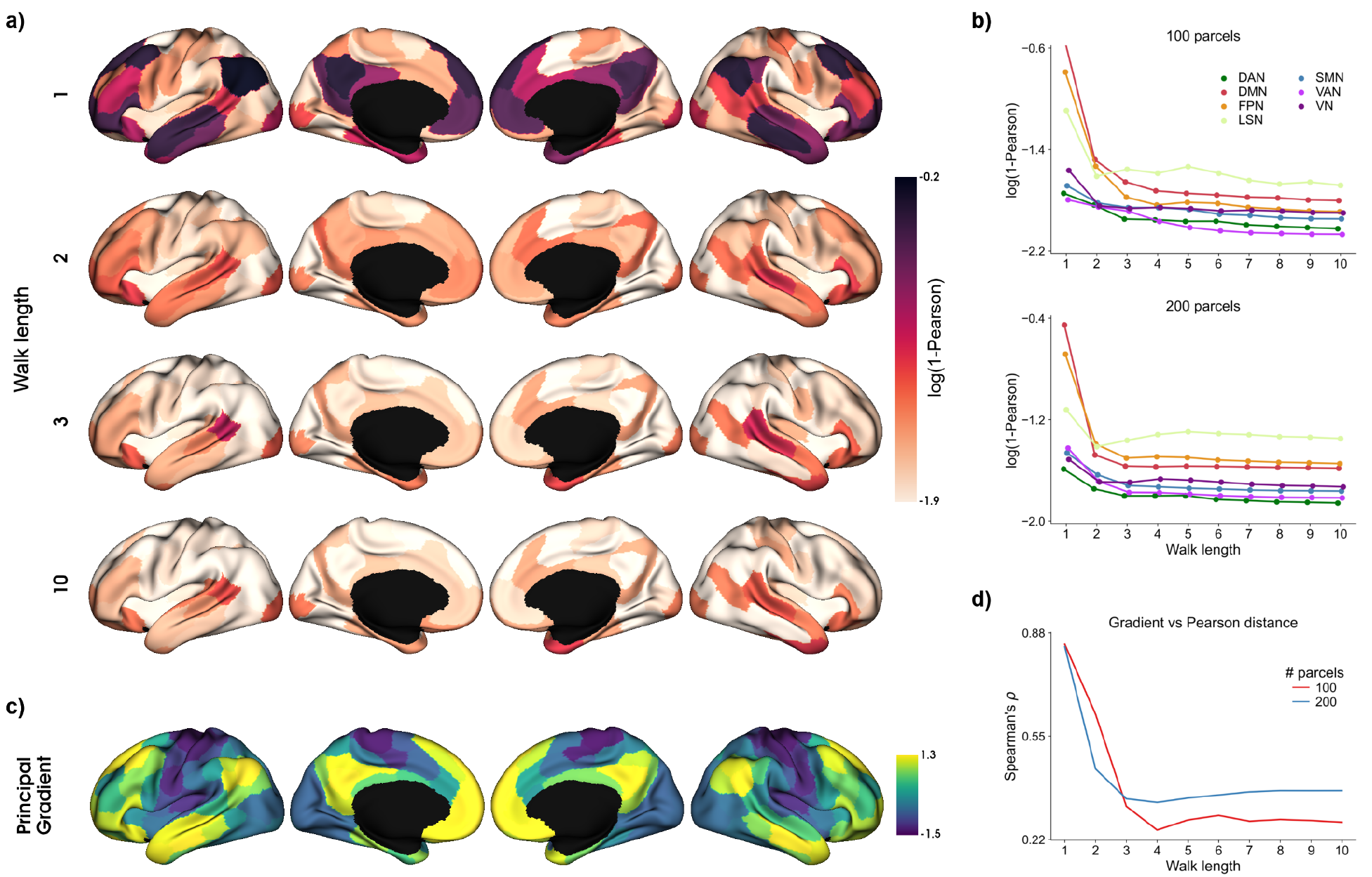
Prediction accuracy at the network level. **a)** Lateral and medial views of left and right hemispheres showing spatial cortical maps of average FC prediction error in terms of log Pearson distance (defined as 1-Pearson) based on the 200-node functional parcellation and the 250 subjects in the holdout data from HCP. Prediction error maps are shown for random walks of length 1, 2, 3 and 10. Each parcel depicts the mean prediction errors across all its edges, with darker colors denoting higher prediction errors. **b)** Network-wise average prediction errors estimated between empirical functional connectivity profiles and the predicted profiles for walks of lengths ranging from 1 to 10. Results are shown for each of the 7 functional Yeo networks based on the 100- and 200-node functional parcellations. **c)** Principal functional gradient derived from the empirical FC in the holdout set of HCP. **d)** Spearman’s correlation of the prediction error maps produced from each walk length and the principal functional gradient. Results are shown for both 100- and 200-node functional parcellations. Abbreviations: dorsal attention (DAN), frontoparietal (FPN), default mode (DMN), visual (VN), limbic (LSN), somatomotor (SMN), and ventral attention (VAN) networks.

### Comparison with state-of-the-art methods

We compared the prediction accuracy (average Pearson’s correlation coefficient) of our proposed approach to several state-of-the-art methods and to other 2 “reduced” versions of our approach (*SingleLength*, *SharedRot*). Benchmarking results are reported in **Table 1** for the experiments in the HCP dataset, and **Table S1** for MICs. These results correspond to using random walks of length 10. In both tables, we found a clear dichotomy in performance between approaches based on the eigenvectors of the SC and the rest (*i.e*., SDK and NLSA). The latter achieved the lowest prediction accuracies, with SDK slightly outperforming NLSA in most experimental settings. It is worth noting that the SDK approach only uses one single diffusion kernel. **Table 1** further reports the performances of the MKL and Spectral approaches, which achieved higher performances than SDK and NLSA, with the Spectral approach producing better predictions than MKL in most scenarios (except when using the 100-node structural parcellation in MICS and the holdout dataset in HCP). Nonetheless, all these approaches were considerably outperformed by our proposed approach in all experimental scenarios, and in both HCP and MICs datasets.

We further assessed the contribution of the different components of our proposed method. As shown in **Table 1** and **Table S1** for HCP and MICs, respectively, the reduced versions of our approach achieved lower prediction accuracies than the original version. When only considering random walks with a single length (*SingleLength*), predictions were more accurate than when using a single rotation shared by all random walks (*SharedRot*). These results underscore the importance of considering random walks of multiple lengths and emphasizes the role of using a different rotation for each length.

Regarding the experimental settings, the functionally-derived parcellation showed to be more beneficial for the prediction of FC than the structural parcellation used in our work. Although the structural parcellation consistently outperformed the functional one with SDK and NLSA, there was a substantial improvement with the functional parcellation over the structural one when considering the best performing approaches (Spectral, SingleLength, SharedRot, Proposed). Finally, we found a clear drop in prediction accuracy across all the methods we evaluated as the number of parcels increased from 100 to 200, regardless of parcellation type (functional or structural) and dataset.

## DISCUSSION

This work presented a Riemannian approach to predict functional connectivity (FC) from the underlying structural connectome (SC) at the individual participant level. The proposed approach leveraged the diffusion maps framework to model the exchange of information through the fibers between different brain regions and learn intermediate kernels capturing this information flow. By capitalizing on manifold optimization, we did not only consider the relationships between structural and functional spectra, but also incorporated a mapping (rotation matrices) between the eigenvectors of both domains. With the proposed approach, we were able to find a robust mapping between the structural and functional embeddings, as shown by our results across different datasets, cortical parcellation atlases and parcellation granularities. Furthermore, our approach allowed us to investigate and understand how brain function gradually emanates from structure as information is propagated through increasingly longer (*i.e*., multiple hops) paths across the structural backbone.

FC does not simply reflect the static and direct wiring of the SC, but it also captures higher-order interactions between potentially only indirectly connected areas (Honey et al., 2009). We hypothesized that by accounting for polysynaptic communication mechanisms, we could explain, to a large extent, the FC observed between pairs of brain regions that lack a direct structural connection. Polysynaptic signaling was modelled by controlling the diffusion time parameter of the diffusion maps framework (Coifman et al., 2005; Coifman & Lafon, 2006). By increasing the diffusion time for a given structural embedding, we were able to generate diffusion coordinates that increasingly captured the interactions between brain regions that were only indirectly connected by structural links. In other words, we were gradually incorporating indirect paths of longer and longer lengths (paths of length 2, 3, 4, and so forth), hence allowing information to flow through increasingly higher-order connections. These interactions were then represented using a radial basis function kernel for each diffusion time. Furthermore, these kernels can be interpreted as intermediate states of the brain while neural information is propagated through the structural fibers. Finally, the different kernels were then fused to provide the predicted FC. Our results showed that, although prediction accuracy increased almost monotonically with increasing diffusion time, there was a clear leveling off around random walks of lengths 3-4, after which only negligible improvements were found. Prior work in the prediction of FC from SC also highlighted shorter structural walks as the strongest contributors to the resulting FC (Becker et al., 2018). This is in line with the economically-optimized configuration of the brain (Bullmore & Sporns, 2012). In addition to minimizing the axonal wiring costs to build the connectome, the expensive metabolic costs spent on information processing also encourage the transfer of information through more economic pathways (Achard & Bullmore, 2007; Avena-Koenigsberger et al., 2014; Bullmore & Sporns, 2012; Laughlin et al., 1998), and hence might be favouring shorter polysynaptic paths.

With diffusion maps, information flow is driven by the transition probabilities of the diffusion operator, which are in turn derived from the edge weights of the original SC matrices. In the current work, we sought to explore the role of different weighting schemes in explaining ongoing brain function: *i)* binary weights only indicating presence or absence of connections, *ii)* weights based on mean streamline length, and *iii)* weights denoting fiber density (conventional scheme). Note that we did not change the connections of the structural graph (*i.e*., add and/or remove edges), only their weights. These weights play a crucial role in guiding the random walker-based information propagation process through the SC. In the binary weighting scheme, the probability of transitioning from one brain region to another is evenly distributed among all its adjacent regions (*i.e*., those with a direct structural connection), whereas in the other weighting schemes, the likelihood of following a particular (direct) connection depends on its length or fiber density. With inadequate weight assignments, we might be misleading the diffusion process and deviating from the true putative flow of information, which can be especially harmful for neural function synchronization of brain regions relying on polysynaptic communication mechanisms. Indeed, this is what we observed with the binary and length-based weighting schemes, which were substantially outperformed when using weights based on fiber density. More importantly, increasing diffusion time showed no consistent contribution to prediction accuracy, and only minor changes were found from one diffusion time to another. Moreover, the length-based weighting scheme attained the lowest prediction accuracy, suggesting that the length of the streamlines might not be the most important indicator of their degree of participation in polysynaptic communication. According to our results, this role appears to be better explained by the density of the streamlines. Regardless of the weighting scheme, our approach relies on diffusion-based communication processes (Coifman et al., 2005; Coifman & Lafon, 2006; Masuda et al., 2017), in which the random walker is only driven by local information. This communication strategy lies on a continuous spectrum of communication processes that ranges from unbiased random walks (such as our proposed approach) at one extreme, through biased random walks that incorporate both local properties and information about the global topology of the structural network, to shortest paths walkers that only consider global information at the other extreme (Avena-Koenigsberger et al., 2014, 2019). Network communication models incorporating both local and global network properties may therefore provide additional information for understanding the correspondence of brain structure and function and hence enhance the predictive power of the proposed approach. Given that signal propagation is strongly influenced by the number of streamlines and also (to a lesser extent) by their lengths (Hermundstad et al., 2013), future work may consider incorporating global information about these properties into the diffusion process to favor transmission of neural signals through shorter and more reliable pathways (Fornito et al., 2016; Goñi et al., 2014).

With the purpose of supporting the dynamic emergence of coherent neural activity patterns, the anatomical substrate of the brain establishes the routing network architecture that facilitates communication between disparate cerebral regions. These structural communication channels have an important say in how brain function is shaped. It is well known that the structure-function relationship is spatially-varying across the cortex, with strong coupling in primary sensory areas that gradually weakens as we move in the sensory-fugal direction (Baum et al., 2020; Suárez et al., 2020; Sydnor et al., 2021; Vázquez-Rodríguez et al., 2019), following a cortical hierarchy of functional and structural organization (Margulies et al., 2016; Paquola et al., 2019; Park et al., 2020). In addition to corroborating these observations, our results showed a tightening of the structure-function coupling in transmodal cortices as diffusion time increased, which, in turn, also translated into a divergence of this coupling from the principal functional gradient. As mentioned above, most of these changes occurred during the first 3 or 4 diffusion timesteps. Although we found some improvements in prediction accuracy with increasing diffusion time in unimodal areas, diffusion time had disproportionately more impact in explaining brain function in transmodal regions. These findings suggest that brain function emerges through polysynaptic communication mechanisms in transmodal cortices, while shorter communication pathways (*e.g*., monosynaptic) are needed in unimodal regions. In a recent study characterizing the directionality of neural information propagation from undirected structural connectome data (Seguin et al., 2019), unimodal and transmodal cortices were found to be at extremes of the send-receive asymmetry spectrum, with unimodal cortices being more likely to be senders and transmodal cortices more likely to be receivers. This communication asymmetry may therefore account for the larger diffusion times required by transmodal cortices, which are thought to integrate multiple streams of information originating from the unimodal sensory areas that require fewer diffusion timesteps.

Most work studying structure-function coupling in the brain investigated this relationship based on group-level SC and FC matrices (Goñi et al., 2014; Seguin et al., 2020; Suárez et al., 2020). Predictions at the group level, however, do not take into account inter-subject variability, *e.g*., using group consensus structural matrices that only consider connections when they are present in at least one-fourth of the participants (Avena-Koenigsberger et al., 2014; Rosenthal et al., 2018). At the individual level, this may translate into less accurate predictions. For example, (Sarwar et al., 2021) using deep learning achieved very high prediction accuracies at the group level (r=0.900) that substantially dropped when considering individual predictions (r=0.550). In this work, our proposed approach was able to accurately predict brain function at the individual level in both HCP and MICs datasets. Across the different parcellation schemes considered, we observed a consistent increase in prediction accuracy when reducing the spatial scale of the parcellations. The type of information used to create the parcellations also had an important impact on out-of-sample performance, with the functionally-derived parcellations typically producing better predictions than the structurally-defined parcellations. In cross-validation, the highest prediction accuracy (r=0.802) was achieved when using the coarsest functional parcellation scheme, whereas the lowest performance (r=0.715) was obtained with the most granular (*i.e*., 200 regions) structural parcellation, both in HCP and MICs. In comparison with the state of the art, our approach attained the best prediction accuracies, regardless of the type of parcellation and dataset. The proposed approach showed solid improvements over the currently best performing method (*i.e*., Spectral), with percentage increases ranging from 2.95% to 14.20% in HCP (across both cross-validation and holdout datasets), and from 11.77% to 18.15% in the MICs dataset. In addition, the ablation analysis of our method highlighted the importance of including different walk lengths and considering multiple rotations (one rotation per length) for providing accurate prediction. In the MICs dataset, for the 200-node structural parcellation, for example, the mean Pearson’s correlation coefficient dropped by 4.63% when only considering a single diffusion time (and hence a single kernel), with an even more substantial drop of 7.30% when using one single rotation shared by all diffusion times. In HCP, prediction accuracy decreased even further with these two reduced versions of our proposed approach. On the other hand, according to our results, the Riemannian regularization (encouraging consecutive diffusion times to use similar rotation matrices) that we initially included in our approach did not show an important contribution to the prediction of FC. This regularization was used to encourage consecutive diffusion times to use similar rotation matrices, however, our results showed that the kernels that contributed the most to the prediction of brain function had very dissimilar rotation matrices, whereas the remaining kernels that barely improved the quality of our predictions had similar rotations. This dichotomy may explain the little contribution of the Riemannian regularization term to the final predictions.

In conclusion, our results show that only a few kernels are necessary to reliably reconstruct functional connectivity from a model of the structural connectome. Moreover, our results underscored that visual, somatomotor and attention networks require generally shorter communication paths than transmodal systems such as the default mode, frontoparietal and limbic networks. The requirement of larger diffusion times in these networks highlights the reliance on more polysynaptic communication mechanisms as we go up the putative cortical hierarchy. Finally, the proposed approach produced highly competitive predictions vis-a-vis current state-of-the-art methods, and this performance improvement was observed across different experimental settings (*i.e*., across different datasets, parcellation schemes, and parcellation granularities). Overall, our findings support a likely contribution of polysynaptic signaling in macroscale brain function, especially in transmodal cortices and thus outline potential mechanisms underlying gradients of structure-function coupling in human cortical networks.

## ACKNOWLEDGEMENTS

Oualid Benkarim was funded by a Healthy Brains for Healthy Lives (HBHL) postdoctoral fellowship and the Quebec Autism Research Training (QART) program. Casey Paquola was funded through a postdoctoral fellowship of the Fonds de la Recherche due Quebec - Santé (FRQ-S). Bo-yong Park was funded by the National Research Foundation of Korea (NRF-2021R1F1A1052303), Institute for Information and Communications Technology Planning and Evaluation (IITP) funded by the Korea Government (MSIT) (2020-0-01389, Artificial Intelligence Convergence Research Center, Inha University; 2021-0-02068, Artificial Intelligence Innovation Hub), and Institute for Basic Science (IBS-R015-D1). Boris Bernhardt acknowledges research support from the National Science and Engineering Research Council of Canada (NSERC Discovery-1304413), the Canadian Institutes of Health Research (CIHR FDN-154298, PJT-174995), SickKids Foundation (NI17-039), Azrieli Center for Autism Research (ACAR-TACC), BrainCanada (Azrieli Future Leaders), and the Tier-2 Canada Research Chairs program. Jessica Royer was funded by a CIHR fellowship. Reinder Vos de Wael was funded by a studentship from the Savoy Foundation. We would also like to acknowledge support from the Helmholtz Foundation and the Healthy Brains for Healthy Lives initiative.

## AUTHOR CONTRIBUTIONS

O.B. and B.C.B. designed the experiments, analyzed the data, and wrote the manuscript. C.P., and B.P. aided with the experiments. J.R., R.R-C., R.V., B.M. and G.P. reviewed the manuscript. O.B. and B.C.B. are the corresponding authors of this work and have responsibility for the integrity of the data analysis.

## CODE AND DATA AVAILABILITY

The code for the proposed framework is publicly available at https://github.com/MICA-MNI/micaopen/sf_prediction. The HCP and MICs dMRI and rs-fMRI data are publicly available at https://www.humanconnectome.org/ and https://portal.conp.ca/dataset?id=projects/mica-mics.

---

**Figure S1.**
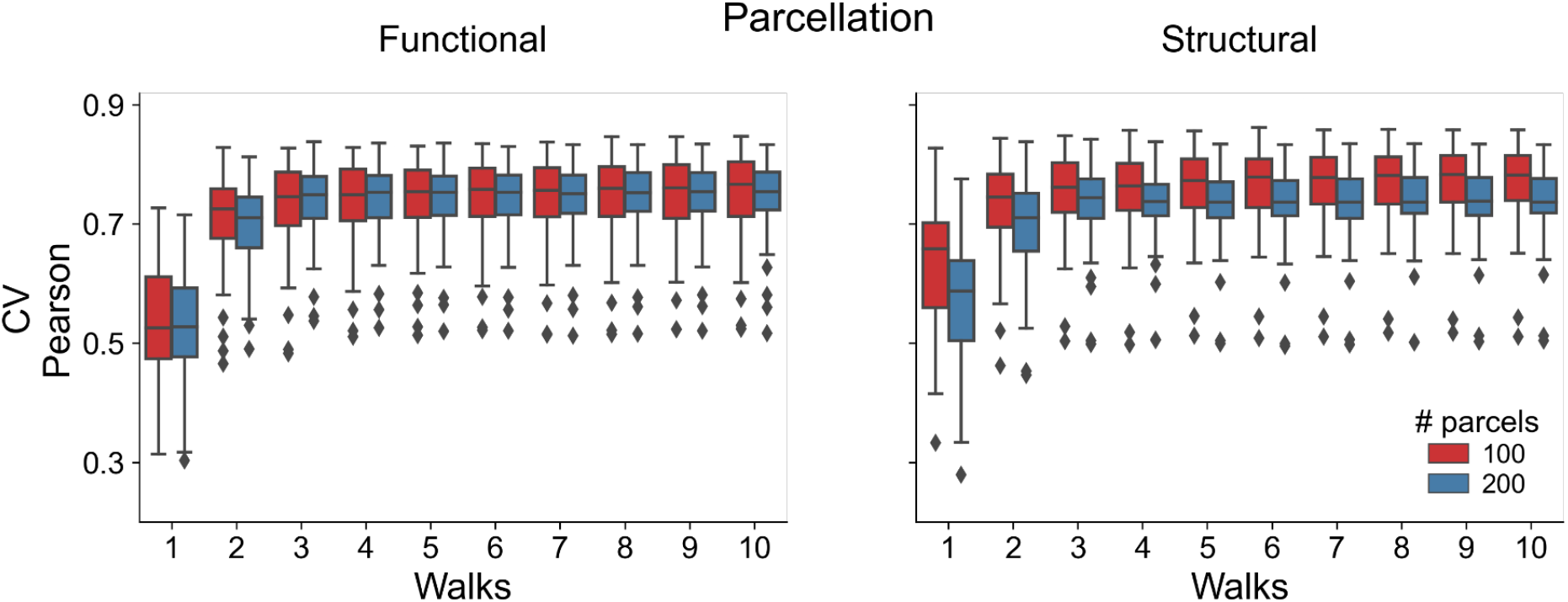
MICs dataset. Boxplots of correlation between empirical and estimated FC matrices using 1 to 10 random walks. Results are reported in cross-validation (top) and holdout (bottom) using functionally- (left) and structurally-derived cortical parcellation (right) with 100 and 200 parcels.

**Figure S2.**
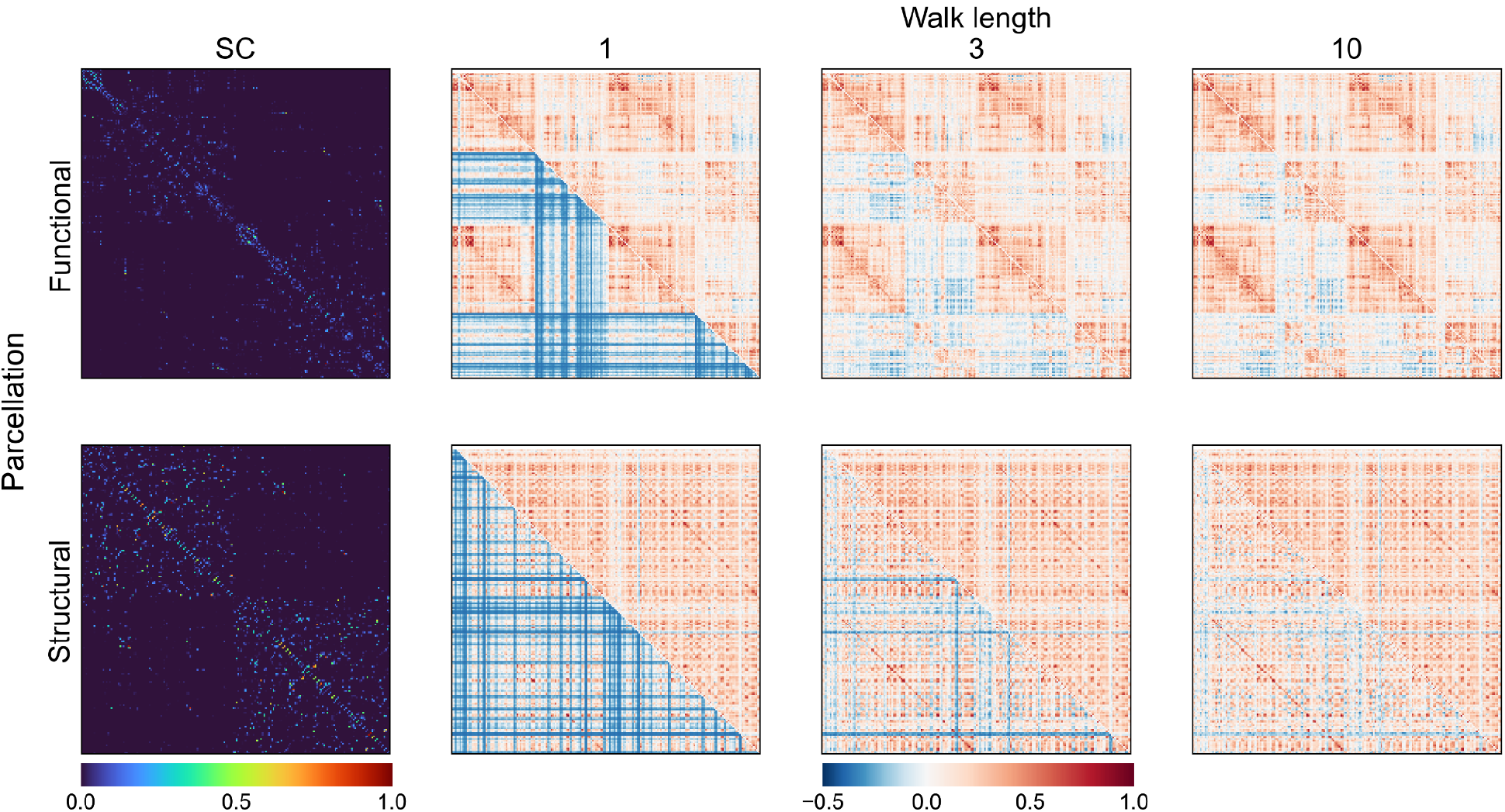
Functional connectivity prediction using random walks of different lengths in the structural connectome in the HCP dataset. Estimated functional connectivity matrices corresponding to the subject that achieved the median Pearson’s correlation coefficient based on functional (top) and structural (bottom) parcellations with 200 parcels. From left to right: structural connectome and estimated FC matrices for random walks of length 1, 3 and 10. Empirical and estimated functional connectivity matrices are shown in upper and lower triangular parts, respectively.

**Table S1.** Mean correlation and standard deviation using SDK, NLSA, MKL, Spectral and the 3 versions of our proposed approach. Performance is reported for 100 and 200 parcels using 3-fold cross validation (CV) in HCP and MICS datasets, and holdout in HCP. For Spectral and our approach, the performance reported corresponds to using 10 random walks, and 16 for MKL.

|  |  | Functional | Structural |
| --- | --- | --- | --- |
| **100** | **SDK** | 0.320±0.045 | 0.427±0.067 |
|  | **NLSA** | 0.299±0.043 | 0.416±0.065 |
|  | **MKL** | 0.632±0.088 | 0.678±0.090 |
|  | **Spectral** | 0.671±0.085 | 0.650±0.081 |
|  | **SingleLength** | 0.723±0.073 | 0.741±0.075 |
|  | **SharedRot** | 0.711±0.070 | 0.734±0.070 |
|  | **Proposed** | **0.750±0.075** | **0.768±0.072** |
| **200** | **SDK** | 0.317±0.042 | 0.338±0.048 |
|  | **NLSA** | 0.303±0.042 | 0.326±0.059 |
|  | **MKL** | 0.601±0.081 | 0.614±0.090 |
|  | **Spectral** | 0.655±0.076 | 0.624±0.079 |
|  | **SingleLength** | 0.727±0.067 | 0.706±0.065 |
|  | **SharedRot** | 0.693±0.069 | 0.685±0.075 |
|  | **Proposed** | **0.742±0.069** | **0.735±0.067** |

